# Temperatures above 37°C increase virulence of a convergent *Klebsiella pneumoniae* sequence type 307 strain

**DOI:** 10.1101/2024.02.06.579160

**Authors:** Justus U. Müller, Michael Schwabe, Lena-Sophie Swiatek, Stefan E. Heiden, Rabea Schlüter, Max Sittner, Jürgen A. Bohnert, Karsten Becker, Evgeny A. Idelevich, Sebastian Guenther, Elias Eger, Katharina Schaufler

**Affiliations:** Department of Epidemiology and Ecology of Antimicrobial Resistance, Helmholtz Institute for One Health, Helmholtz Centre for Infection Research HZI, Greifswald, Germany; Imaging Center of the Department of Biology, University of Greifswald, Greifswald, Germany; Friedrich Loeffler-Institute of Medical Microbiology, University Medicine Greifswald, Greifswald, Germany; Institute of Medical Microbiology, University Hospital Münster, Münster, Germany; Pharmaceutical Biology, Institute of Pharmacy, University of Greifswald, Greifswald, Germany; University Medicine Greifswald, Greifswald, Germany

**Keywords:** *K. pneumoniae*, temperature-dependent virulence, hypermucoviscosity, hypervirulence, plasmid copy number, transcriptomics

## Abstract

Hypermucoviscosity in *Klebsiella pneumoniae* is often related to the overexpression of capsular polysaccharides, regulated by complex biosynthetic mechanisms in response to external cues. However, little is known about the processes involved in hypermucoviscosity in convergent *K. pneumoniae*, which combine extensive drug resistance with high bacterial virulence, under pathophysiological conditions. This study aimed to fill this gap by investigating the temperature dependence of hypermucoviscosity and overall virulence in a convergent *K. pneumoniae* strain isolated during a clonal outbreak belonging to the high-risk sequence type (ST)307.

Hypermucoviscosity, biofilm formation, and mortality rates in *Galleria mellonella* larvae were examined at different temperatures (room temperature, 28°C, 37°C, 40°C and 42°C) and with various phenotypic experiments including electron microscopy. The underlying mechanisms of the phenotypic changes were explored via qPCR analysis to evaluate plasmid copy numbers, and transcriptomics.

Our results indicate a temperature-dependent “switch” above 37°C to a hypermucoviscous phenotype, correlating with increased biofilm formation capacity and *in vivo* mortality, which might be due to a bacterial response to pathophysiological conditions, i.e., fever. In addition, we detected upregulation of a hybrid plasmid encoding both carbapenemase and the mucoid regulator *rmpA* genes. Surprisingly, *rmpA* did not exhibit temperature-dependent differential gene expression, suggesting other drivers. Apparent co-regulation of hypermucoviscosity and fimbrial expression was also identified.

This study not only revealed the impact that increased temperatures above 37°C have on hypermucoviscosity and virulence in a convergent *K. pneumoniae* strain but contributes to the understanding of previously unrecognized dimension of *K. pneumoniaés* behavior, emphasizing its adaptability to changing environments.

**Abstract importance:** Understanding the temperature-dependent dynamics of hypermucoviscosity in *Klebsiella pneumoniae* is crucial for unraveling the intricacies of its hypervirulence. This study investigates a convergent *K. pneumoniae* strain, ST307, revealing a temperature-dependent switch to hypermucoviscosity above 37 °C. The findings showcase a correlation between increased temperature, hypermucoviscosity, enhanced attachment, and heightened *in vivo* mortality. Notably, a hybrid plasmid encoding carbapenemase and mucoid regulator genes was upregulated at elevated temperatures. The study sheds light on previously unexplored aspects of *K. pneumoniae* behavior, emphasizing its adaptability in response to changing environments. The identified temperature-associated regulatory mechanisms offer insights into the pathogen’s response to fever, contributing to our broader understanding of bacterial adaptation. This research contributes to addressing the global challenge of hypervirulent, drug-resistant *K. pneumoniae* strains, providing valuable implications for future treatment strategies.

## 1. Introduction

The opportunistic pathogen *Klebsiella pneumoniae* is frequently associated with nosocomial infections worldwide including pneumonia, urinary tract, and bloodstream infections [1]. Classic, mostly nosocomial *K. pneumoniae* (cKp) often affects individuals with compromised immune systems [2] exacerbated by the rise of multidrug-resistant (MDR) representatives [3]. Beyond the hospital walls, hypervirulent, *K. pneumoniae* (hvKp) strains can cause infections in healthy individuals [4]. HvKp harbors a repertoire of key virulence factors such as siderophores [5]. In recent years, the global emergence of converging pathotypes of *K. pneumoniae* strains contributed to difficult-to-treat infections, as they combine extensive drug resistance with hypervirulence mostly driven by hybrid plasmids [6]. Traditionally, *K. pneumoniae* hypervirulence has been identified through a positive string test [7]. This test explores “hypermucoviscosity”, a characteristic associated with better evasion of macrophages contributing to the invasive potential of hvKp [8]. Despite the assumption that hypervirulence and hypermucoviscosity are connected, there is evidence that hypermucoviscosity is not a peculiar marker of hypervirulence [9]. Precise determination of hypervirulence involves *in vivo* experiments and specific genetic markers like *iutA* (hydroxamate siderophore), *iroB* (catecholate siderophore), and *rmpA* or *rmpA2* (regulator of the mucoid phenotype) [8]. Hypermucoviscosity correlates with clinical outcomes such as pyogenic liver abscesses [10]. While more than 79 different *Klebsiella* capsule types exist, the various capsule types seemingly differ in the composition of the capsule, which influences virulence and hypermucoviscosity. Especially monosaccharides such as mannose and rhamnose seem to play a role [11]. Other complex biosynthetic and regulatory mechanisms responding to external stimuli such as iron availability and carbon sources are also involved [12]. While mechanisms for bacterial adaptation to host temperatures are well-established, the impact of temperature changes within the host, such as fever episodes, on the regulation of virulence factors is not fully explored. Notably, there is a significant gap in understanding the temperature effects on hypermucoviscosity of *K. pneumoniae*, particularly above 37 °C [13, 14].

Here, we investigated capsule production and hypermucoviscosity in a convergent *K. pneumoniae* ST307 strain at different temperatures. By combining omics with *in vitro* and *in vivo* phenotypic experiments, we revealed temperature dependence of hypermucoviscosity and additional bacterial virulence, which are seemingly based on plasmid copy number (PCN)- and transcriptional changes.

## 2. Results

We investigated hypermucoviscosity and mortality for a previously published, convergent *K. pneumoniae* ST307 strain (PBIO1953 [15]) at RT, 28 °C, 37 °C, 40 °C, and 42 °C (Figure 1). First, we stained capsular polysaccharides (Figure 1A). Here, white to yellow-beige colonies indicate “normal” exopolysaccharide production, which applied to PBIO1953 at RT, 28 °C, and 37 °C. In contrast, black colonies appeared above 37 °C (Figure 1A, Figure Appendix (A) 6), implying increased polysaccharide biosynthesis [16]. String tests confirmed these result (data not shown), hypermucoviscosity was identified at 40 °C and 42 °C. The sedimentation assay also confirmed higher viscosity at 40 °C (*p* = 0.0325) and 42 °C (*p* = 0.0187) compared to 37 °C, indicating increased capsular polysaccharide production (Figure 1B). Third, biofilm experiments revealed that increasing temperatures led to higher affinity of PBIO1953 to adhere to abiotic surfaces (Figure 1C). A significant increase in specific biofilm formation was observed at 40 °C (*p* = 0.0053) and 42 °C (*p* = 0.0049) in comparison to 37 °C. Interestingly, this was not related to curli or cellobiose production (Figure (A) 1).

**Figure 1.**
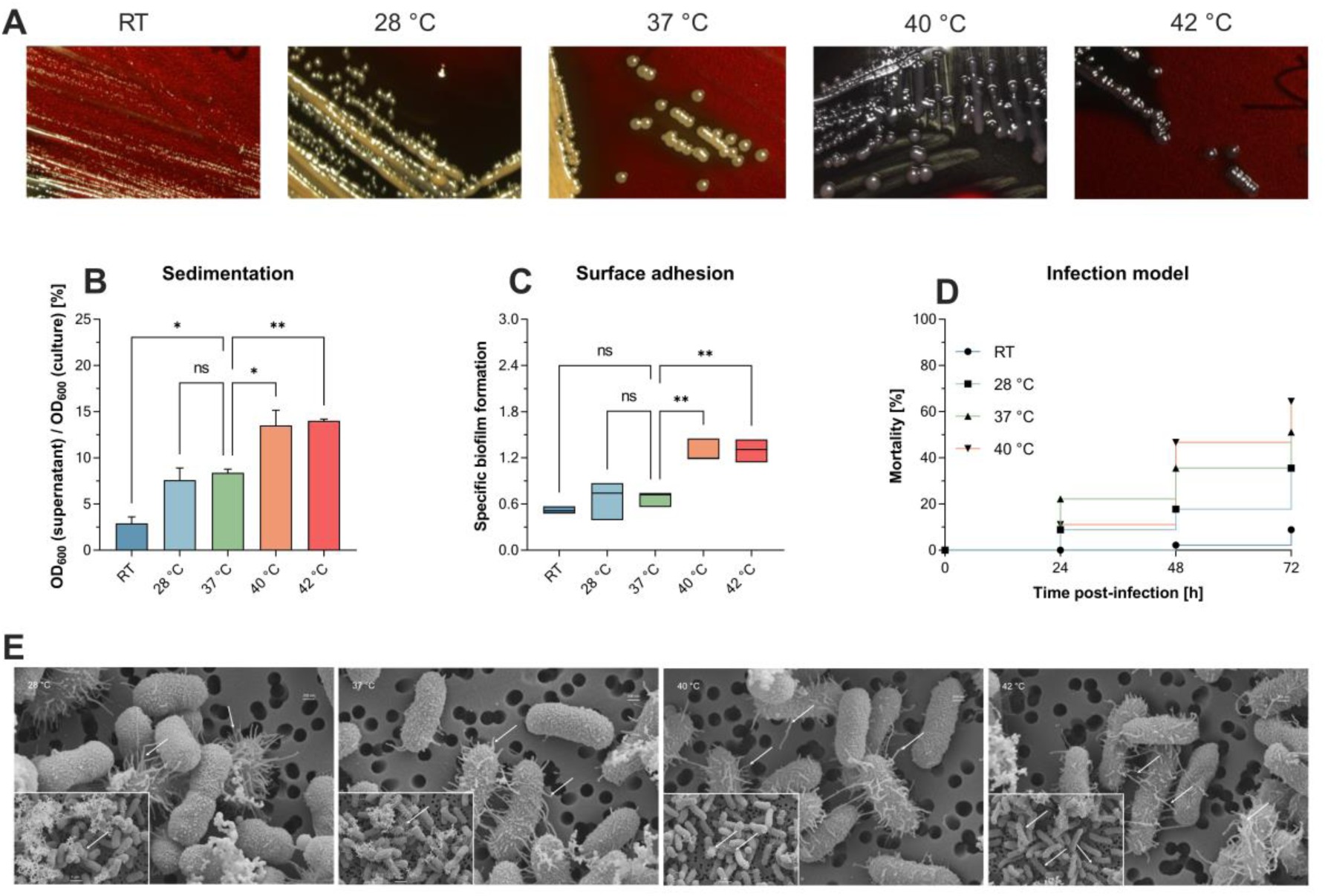
Different temperatures affect mucoviscosity and overall virulence of the convergent *K. pneumoniae* ST307 strain PBIO1953. **A** Staining of capsular polysaccharides revealed a temperature-dependent change from a “normal” mucoid phenotype (yellow-beige colonies) to a hypermucoid phenotype (black colonies) at 40 °C and 42 °C. **B** The increased production of capsule polysaccharides was associated with a decrease in sedimentation upon centrifugation at 1,000 x g for 5 min. Results are shown as mean and standard error of percentage OD_600_ remaining in the supernatant after centrifugation (n = 3). **C** Temperatures above 37 °C resulted in increased adhesion to plastic surfaces. Results are expressed as growth-adjusted specific biofilm formation. The line in the box indicates the median value (n = 3). **D** The in vivo infection model showed a temperature-dependent increase in mortality rates. Kaplan-Meier plot of mortality rates in the *G. mellonella* larvae (n = 30). Results are expressed as mean percent mortality after injection of 2 × 10^5^ CFU/larvae. For all results, mucoviscosity-associated characteristics at the different temperatures were compared to 37 °C using analysis of variance (one-way ANOVA with Dunnett’s multiple comparison post hoc test); ns, not significant; *P*\* <0.05; **, *P* <0.01. RT, room temperature. **E** Scanning electron micrographs of PBIO1953 at 28 °C, 37 °C, 40 °C, and 42 °C at 20,000x magnification, scale bar = 200 nm (inserts: 10,000x magnification, scale bar = 1 µm), arrowheads show fimbriae -like structures.

Finally, to explore the impact of different temperatures on the overall virulence of PBIO1953, we assessed mortality rates in *Galleria (G.) mellonella* larvae (Figure 1D). After 24 hours, mortality rates at 37 °C (22.22%) exceeded those observed at 40 °C (11.12%). However, 48 hours post-infection, mortality rates at 40 °C consistently exceeded those at 37 °C, with a peak after 72 hours. As control, the larvae were mock-infected with PBS and incubated at RT, 28 °C, 37 °C, 40 °C with no detected temperature influence on the larvae mortality (Figure A3). Note that incubation of mock-infected *G. mellonella* larvae at 42 °C resulted in mortality rates greater than 10% (data not shown) (e.g., [17]).

Interestingly, scanning electron micrographs indicate an increasing amount of virulence- and biofilm-associated fimbriae structures with higher temperatures (Figure 1E), reinforcing earlier observations. It is important to note that the bacterial capsule appears compromised during the staining process. Nevertheless, the visible increase of these structures at higher temperatures (40 °C, 42 °C) implies a temperature-associated regulation [18]

### Different temperatures affect PCN

To explore the underlying mechanisms, PCN-variations were measured using qPCR. Previously, we have shown that the convergent PBIO1953 strain harbors five different plasmids, one of which is a hybrid plasmid (plasmid 1) encoding both AMR and hypervirulence genes [15]. The three largest PBIO1953 plasmids were included in the subsequent analysis: plasmid 1 (360,596 bp), positive for the metallo-β-lactamase gene *bla*_NDM-1_ and the regulator of the mucoid phenotype *rmpA*, plasmid 2 (130,131 bp) encoding the extended-spectrum β-lactamase (ESBL) gene *bla*_CTX-M-15_, and plasmid 3 (72,679 bp) without any resistance genes (Figure 2).

**Figure 2.**
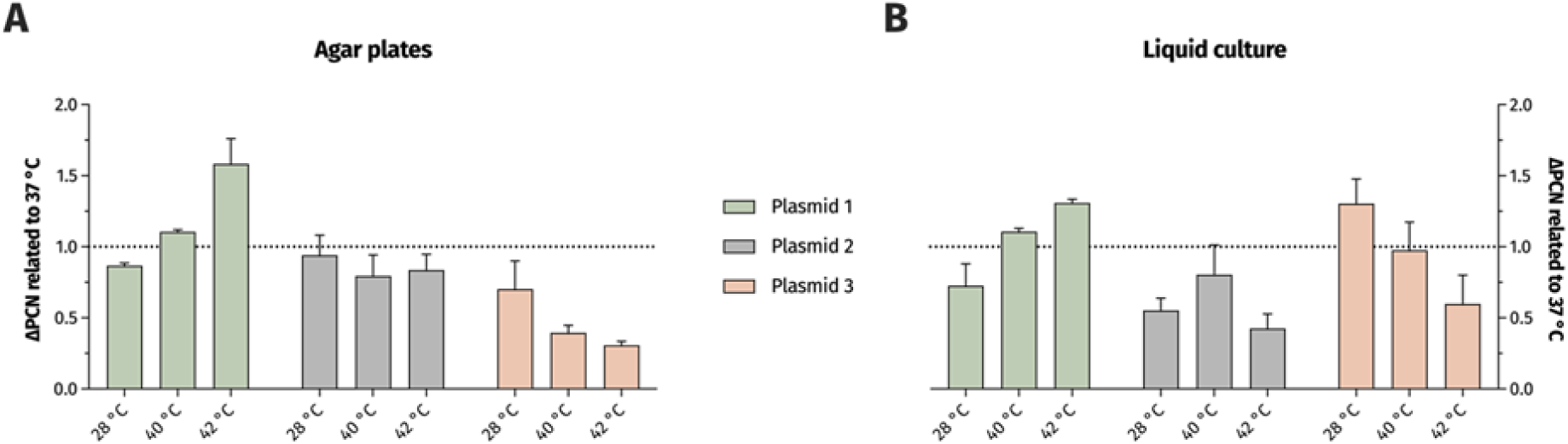
Temperature variation affect PCN. The PCN was determined from individual colonies grown on congo-red-dye-enriched agar plates (**A**) or from cells pelleted from sedimentation assay cultures (**B**). The results are expressed as mean and standard error of the change in the normalized PCN.

As the hypermucoviscosity switch was observed above 37 °C, we normalized all PCN to 37 °C. Interestingly, the PCN differed not only based on the different temperatures but also regarding the respective plasmid and experimental set-ups (Figure 2). For the hybrid plasmid 1, the PCN increased with higher temperatures, with the highest value obtained at 42 °C. The PCN of plasmid 2 did not show a clear temperature decency, but a PCN reduction was apparent at both 28 °C and 42 °C in the liquid culture set-up. Plasmid 3 showed a PCN decreasing with higher temperatures.

### Temperature-dependent transcriptomic changes

For transcriptomics, we first focused on genes displaying differential gene expression (DGE) at 28 °C, 40 °C and 42 °C (e-value < 0.05, |log2fold| change > 1.5) in comparison to 37 °C. DGE appeared mostly between 28 °C and 37 °C (Figure A 2, Figure (A) 4), while, at 37 °C compared to 40 °C and 42 °C, we only noticed few differentially expressed genes. Our initial hypothesis that *rmpA* would be differentially expressed was rejected and, genes related to the capsular gene cluster were unaffected.

When comparing 28 °C-associated gene expression to 37 °C (and 40 °C and 42 °C), we observed various, significant changes. Notably, the chromosomally located *ibpAB* gene and two small heat shock proteins encoded on plasmid 2, exhibited differential expression at increased temperatures. These genes are known for mitigating growth defects under thermal stress [19]. Furthermore, we observed the upregulation of several stress-related genes at 37 °C and higher, including the Fur repressor, which possesses protective properties against reactive oxygen species [20] and *sodA*, a gene involved in radical scavenging. Of particular interest was the upregulation of *acrAB*, known for its role in antimicrobial resistance and virulence [21]. Remarkably, the expression of *mrkABCD*, which encodes the type 3 fimbriae system, increased at 37 °C and above. Previous research has suggested that environmental factors [22], like temperature, can impact attachment factors, making this observation particularly intriguing. Conversely, we noted a downregulation of *fimHGFDCIA,* responsible for the expression of fimbriae type 1 [23], and the downregulation of the *rfb* gene, which encodes O-antigens [24]. Additionally, *ompA*, a known virulence factor and key element in immune system recognition [25] was downregulated at increased temperatures. On plasmid 1, the partitioning system *parAB* responsible for segregation and plasmid stability [26] demonstrated increased expression at higher temperatures. Notably, genes associated with transposon activation and IS sequences were also upregulated on plasmid 1. The gene *iutA,* encoding aerobactin and contributing to bacterial virulence, was upregulated at 37 °C (and 40 °C and 42 °C). Moreover, on plasmid 2, the metallo-resistance gene *arsR* was upregulated [27]. Finally, plasmids 4 and 5 showed increased expression of genes involved in conjugative transfer, including *traA* [28]and *mobC* [29]. When comparing the transcriptomes of 40 °C to those at 37 °C, DGE was observed in six genes. Notably, two downregulated genes*, metR* (associated with the methionine pathway) and *argC* (linked to the arginine pathway), were particularly interesting as they are important for general growth. Only two genes were upregulated, with *betB* standing out as this gene is involved in the biosynthesis of osmoprotective choline-glycine betadine [30]. Similarly, the comparison of 42 °C to 37 °C revealed six differentially expressed genes. *betB* demonstrated concordance with the upregulated genes at 40 °C. An intriguing finding was the upregulation of *pecM*, a gene previously implicated as a potential deactivator of the *fim* operon and a member of the permease of the drug/metabolite transporter (DMT) superfamily.

## 3. Discussion

Hypermucoviscosity is often referred to as the most important virulence characteristic of hvKp. However, it is largely unclear how external stressors can affect capsule expression and thus virulence in bacteria, especially in convergent *K. pneumoniae*. We revealed that a convergent *K. pneumoniae* ST307 outbreak strain [15], changed from a “normal” to a hypermucoid and pronounced virulent phenotype upon temperature exposure above the healthy human body temperature at about 37 °C.

Bacterial capsule formation is a highly complex process influenced by several internal and external factors. Although hvKp frequently exhibits capsule types K1 and K2, a clear correlation between the capsule type and hypervirulence has not yet been established [31]. To our knowledge, we are the first to report a temperature-dependent phenotypic switch for *K. pneumoniae*. Previously, Le et al. (2022) have shown that temperatures below 37 °C may affect hypermucoviscosity, based on the upregulation of plasmid-encoded *rmpA* at 37 °C when compared to RT [32]. In contrast, our results suggest that *rmpA*, although also plasmid-encoded, was not differentially expressed at different temperatures. More importantly, the increased PCN plasmid 1 does seemingly not have any effect on *rmpA* expression, although it is known that higher gene abundance as a result of increased PCN can lead to higher gene expression. It seems likely that hypermucoviscosity does not exclusively depend on *rmpA* expression. This assumption has been previously supported by studies where *K. pneumoniae* isolates harbor impaired capsule locus genes and the respective regulator but show hypermucoviscosity [33].

While a substantial number of genes exhibited DGE between 28 °C and 37 °C, only a limited number of genes was differentially regulated at 37 °C vs. 40 °C or 42 °C. For all comparisons, most of the differentially expressed genes were chromosomally encoded, which suggests that increasing temperatures up to pathophysiological conditions (i.e., fever) does not lead to major shifts in the transcriptome and to the phenotypic switch. Rather, this seems to fine-tune the expression of a small number of genes together with the different PCN. We surmise that temperature-dependent stress regulators and heat shock proteins may be involved in the regulation of capsule production, which is supported by previous studies [34]. However, why the upregulation of stress response regulators and proteins, which normally control the expression of a large number of protein-associated genes [19], does not lead to a clearer change in the bacterial transcriptome, remains unclear. The only direct regulatory pathway previously connected to capsule synthesis and influenced by increased temperatures was the *rcsA* gene, which contributes to capsular overexpression and is usually expressed at temperatures below 37 °C [35]. Note that this is contrary to our findings.

Here, we identified a link between hypermucoviscosity and fimbrial regulation. The upregulation of type 3 fimbriae at temperatures above 28 °C might be responsible for the fimbrial-like structures at higher temperatures as visualized by scanning electron microscopy. Interestingly, Lin et al. (2011) showed that hypermucoviscosity and type 3 fimbriae production can be co-regulated and depend on iron availability, and that iron limitation results in a reduction of mucoviscosity and fimbriae expression [36]. Conversely, we observed a downregulation of type 1 fimbriae at higher temperatures. This might be because type 1 and type 3 fimbriae are independently regulated. However, in the CV assay, which measures specific biofilm formation, increased attachment at 40 °C (and 42 °C) compared to 37 °C was measured, indicating a higher cell-to-cell and cell-to-plastic adherence, which could be fimbriae type 3-related and is essential in the transition from a planktonic to a multicellular lifestyle.

We speculate that PBIO1953 has evolved specific mechanisms to respond to the host’s immune system, combined with a temperature-dependent formation of a physical capsule barrier and increased attachment at 40 °C and 42 °C. In the *in vivo* infection model, higher mortality was initially achieved after 24 hours at 37 °C than at 40 °C, which may be due to the initial encapsulation process combined with higher adherence at 40 °C. As a result, the immune system response may be impeded, enabling bacterial proliferation. After 48 h and 72 h, we noticed a subsequent mortality increase at 40 °C, again hinting towards better bacterial proliferation, possibly due to encapsulation and the previously mentioned changes. Similar has been previously suggested for various pathogens [37] but not *Klebsiella*. At the same time, we must acknowledge that some of our results are based on RNA sequencing data with stringent cutoffs and obtained from liquid cultures, which adds some uncertainty when trying to apply these findings to *in vivo* situations. A further limitation is the fact that we only investigated a single *K. pneumoniae* strain. Prospective studies will have to explore whether similar applies to other strains. Nevertheless, our results suggest that enhanced temperatures and thus pathological conditions (i.e., fever) can lead to increased bacterial virulence, further exacerbating the overall tense situation. However, hypermucoviscosity could also represent a vulnerability that could be considered in a future treatment strategy.

## 4. Conclusions

Here we show that increased temperatures and thus potentially pathological conditions led to increased virulence in a convergent *K. pneumoniae* strain, which might play a pivotal role in shaping the dynamics of infection processes. In addition, our study contributes to a better understanding of the underlying mechanisms leading to hypermucoviscosity and *in vivo* bacterial virulence.

## 5. Methods & Methods

### Bacterial strains

| Name | Species, source |
| --- | --- |
| PBIO1953 | *K. pneumoniae*, [15] |

### Hypermucoviscosity

In the string test, a sterile loop was applied to a single colony on an agar plate, which had been cultured at varying temperatures (28 °C, 37 °C, 40 °C, 42 °C) for a duration of 24 h. The loop was gently lifted, and if there was a 5 mm extension without breaking, the string test was considered positive, which characterizes hypermucoviscosity [38]. Furthermore, the hypermucoviscosity was confirmed with the sedimentation assay as described previously [12]. In short, 50 µL of a standardized bacterial suspension (0.5 McFarland standard turbidity in 0.9 % (w/v) aqueous NaCl solution) was transferred into 5 mL LB broth and was incubated at the desired temperature (RT, 28 °C, 37 °C, 40 °C, 42 °C) for 24 h. Afterwards 1.5 mL of the 24 h culture was transferred into a 2 mL reaction tube and centrifuged for 1000 x g for 5 minutes, 200 µL from the supernatant as well as from the 24 h culture were pipetted as triplicates into a 96 well plate and the OD_600_ was measured with a plate reader (CLARIOstarplus, BMG LABTECH, Ortenberg, Germany) [39].

The ability to generate exopolysaccharides was assessed using BHI agar plates supplemented with 5 % sucrose (w/v) and 0.08 % congo-red (w/v), following the methodology outlined in a prior study [16]. A single colony was picked up from an LB agar plate, streaked onto the stained BHI agar plates, and subsequently incubated overnight for 24 h at (RT, 28 °C, 37 °C, 38 °C, 39 °C, 40 °C, 42 °C). If the single colonies color black, they would be classified as positive for exopolysaccharides, while single colonies which are white to yellow would be considered negative for exopolysaccharide production.

### Attachment and specific biofilm formation

The temperature influence on attachment factors was investigated with the crystal violet assay, as described previously [40], [41] In short 50 µL of an overnight culture was transferred to 5 mL LB. After a visible turbidity samples were adjusted to an OD_600_ of 0.01 and 200 µL was transferred as into a 96 well plate. Afterwards, they were incubated at 28 °C, 37 °C, 40 °C and 42 °C for 24 h. Following the incubation, the OD_600_ was measured to determine the overall growth. Then, the supernatant of each well was discarded, the plate was washed three times with deionized water and dried for 10 min. Subsequently, fixed with 250 µL methanol for 15 min and then air dried for 15 min. Now the remaining cells were stained with 250 µL 1 % crystal violet solution (w/v) for 30 min. This was followed by three washing steps with deionized water and drying for 10 min. The stained bacteria were dissolved with 300 µL of an ethanol acetone mixture with a ratio of 80 to 20 (v/v) and shaken for 30 min at 220 rpm at room temperature. Then the OD_570_ was measured. The specific biofilm formation (SBF) [42] was then calculated. The negative control was subtracted from the OD_570_ of the sample and then divided by the OD_600_ of the sample.

### Galleria mellonella in vivo infection model

To test the influence of different temperatures on the pathogenic potential the *Galleria mellonella in vivo* infection model was used, as previously described [43] with slight adjustments of the experiments setup in respect to this temperature dependent study. Briefly, 2 mL of overnight culture of PBIO1953 was harvested and pelleted at 16000 x g for 5 min at RT. The pellet was once washed with PBS and diluted to an OD_600_ of 1.0. The 2 × 10^9^ CFU/mL were further adjusted to 2 × 10^6^ CFU/mL. The larvae (from Peter Zoopalast, Kiel, Germany) were randomly divided into 6 different groups of 10 individuals. 10 µL of the bacterial suspension was injected into the right proleg of the larvae. For the control groups 10 µL of PBS solution was used as injection, to check if the Infection of *Galleria mellonella* larvae was affected by the traumata or the altered incubation temperature. The sample and control groups were incubated in a 90 mm glass petri dish at 28 °C, 37 °C and 40 °C each. The vital state of the larvae was controlled every 24 h. The larvae were considered dead, if they showed strong pigmentation accompanied by not responding to physical stimuli. This assay was performed three independent times the results were pooled for each condition. A Kaplan-Meier plot was generated to visualize the mortality rates.

### Plasmid copy number (PCN)

To determine the plasmid copy numbers of the three largest plasmids (plasmid 1 *bla*_NDM-1_ and *rmpA*, plasmid 2 *bla*_CTX-M-15_, plasmid 3) at different temperatures (RT, 28 °C, 37 °C, 40 °C, 42 °C) qPCR was used. Therefore, sets of different Primers were designed to amplify a specific region on the gene constructs of interest. The primers were designed according to the manufacture’s instruction for Luna qPCR Kit (New England Biolabs GmbH, Ipswich, MA, US) and ordered from Eurofins (Eurofins Genomics Europe Shared Services GmbH, Ebersberg, Germany).

The calibration for the plasmids (1-3) and the chromosome was performed with the Q5 High-Fidelity DNA Polymerase (New England Biolabs GmbH, Ipswich, MA, US) according to the manufacture’s guideline, and isolated with NucleoSpin Gel and PCR Clean-up (MACHEREY-NAGEL GmbH & Co. KG, Düren, Germany) and quantified with the Qubit 4 fluorometer using the dsDNA HS Assay Kit (Thermo Fisher Scientific, Waltham, MA, USA). The qPCR was performed with the Luna qPCR protocol (New England Biolabs GmbH, Ipswich, MA, US) in regard to the qPCR cycler CFX Opus 96 Real-Time PCR System (Bio-Rad, Hercules, CA, US). Each sample was set up as seen in the Table 2 and each sample was measured in triplicates following the protocol shown in Table 3.

**Table 1.**
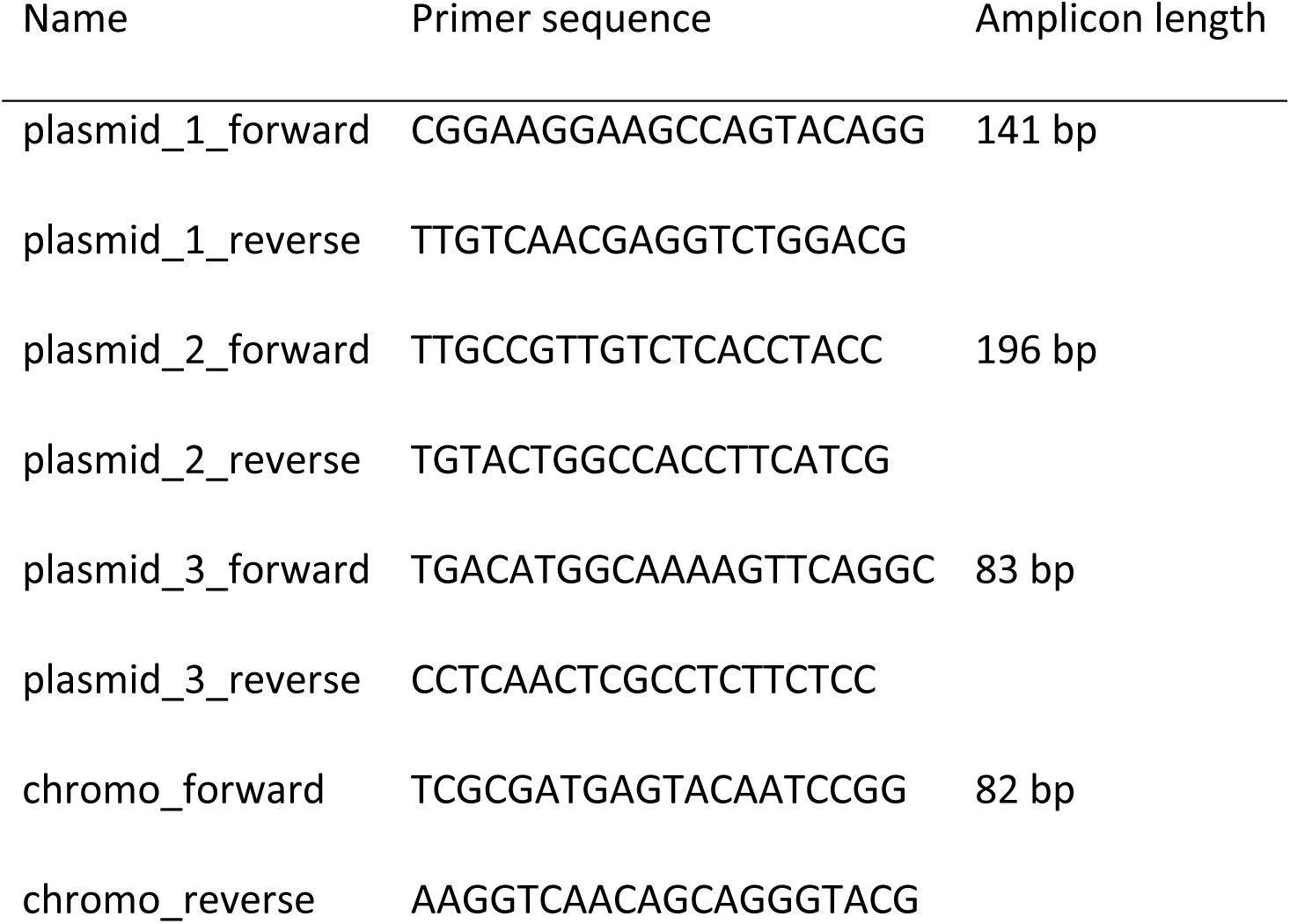
Primer Sequences for plasmid copy number analyses of plasmid 1-3, and chromosome.

**Table 2.** qPCR reaction setup per well.

| Component | 20 µl reaction |
| --- | --- |
| Luna Universal qPCR Master Mix | 10 µl |
| Forward primer (10µM) | 0.5 µl |
| Reverse primer (10µM) | 0.5 µl |
| Template DNA | 50 ng |
| Nuclease free water | to 20µl |

**Table 3.** qPCR thermocycler protocol for the Biorad DFX Optus 96-well, qPCR Luna Mastermix.

| Cycle step | Temperature | Time | Cycles |
| --- | --- | --- | --- |
| Initial Denaturation | 95 °C | 60 s | 1 |
| Denaturation Extension | 95 °C 60 °C | 15 s 30 s (+plate read) | 45 |
| Meltingcurve | 60 °C -> 95 °C | Biorad CFX Optus standard | 1 |

The measured concentration of the samples was related to the overall length of the gene structure and the plasmid copy number was calculated as

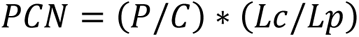

PCN: Plasmid copy number per genome equivalent P: Overall amount of the plasmid (in ng) C: Overall amount of the chromosome (in ng) Lc: Overall length of the chromosome (in base pairs) Lp: Overall length of the plasmid (in base pairs).

### Electron microscopy

A single colony was inoculated into 30 mL of LB in an Erlenmeyer flask and cells were incubated at 28 °C, 37 °C, 40 °C and 42 °C for 24 hours. Subsequently, the bacterial suspension was adjusted to OD_600_ = 1, and 1 mL of the suspension was diluted with 9 mL of 0.9% NaCl. The resulting suspension was then filtered through a hydrophilic polycarbonate filter (0.2 μm pore size, Merck Millipore). A segment of the filter was fixed with 1% glutaraldehyde and 4% paraformaldehyde in washing buffer (10 mM cacodylate buffer, 1 mM CaCl_2_, 0.075% ruthenium red; pH 7) and then the samples were stored at 4 °C until further processing (not more than 16 hours).

Samples were washed with washing buffer three times for 10 min each time , treated with 2% tannic acid in washing buffer for 1 h at RT, and then washed again with washing buffer three times for 15 minutes each time. Afterwards, the samples were dehydrated in a graded series of aqueous ethanol solutions (10, 30, 50, 70, 90, 100%) on ice for 15 min each step. Before the final change to 100% ethanol, samples were allowed to reach RT and then critical point-dried with liquid CO_2_. Finally, the samples were mounted on aluminum stubs, sputtered with gold/palladium, and examined with a field emission scanning electron microscope Supra 40VP (Carl Zeiss Microscopy Deutschland GmbH, Oberkochen, Germany) using the Everhart-Thornley SE detector and the in-lens detector in a 70:30 ratio at an acceleration voltage of 5 kV. All micrographs were edited by using Adobe Photoshop CS6.

### DNA Isolation

The DNA isolation for the qPCR was performed by either 1 mL of 24 h cultures in LB at 28 °C, 37 °C, 40 °C and 42 °C or directly from the BHI agar plates supplemented with congo-red and sucrose incubated at the 4 different temperatures, where 4 to 6 single colonies were scraped off and resuspended via vortexing in 500 µL of PBS in a 1.5 mL test tube. The DNA isolation was then performed with the *MasterPure DNA Purification Kit for Blood* according to the manufacturer’s specifications (Lucigen, Middleton, WI, USA). The quantification of the DNA was ensured by using the Qubit 4 fluorometer using the dsDNA HS Assay Kit (Thermo Fisher Scientific, Waltham, MA, USA).

### RNA Isolation

For transcriptomic analyses the RNA was isolated from 50 mL liquid 24 h cultures at (28 °C, 37 °C, 40 °C and 42 °C). 800 µL of the cultures were harvested in a 1.5 mL reaction tube and instantly chilled with liquid nitrogen for 5 s, to inhibit changes in the transcriptome. After the centrifugation (16,000 × *g* for 3 min at 2 °C), the following RNA isolation was performed immediately with Monarch™ Total RNA Miniprep Kit (New England Biolabs GmbH, Ipswich, MA, US) according to the manufacture’s instruction. The quality of the isolated RNA was tested using the Bioanalyzer with the Total RNA Nano Chip (Agilent, Santa Clara, CA, US).

The samples were shipped on dry ice to Novogene (NOVOGENE COMPANY LTD., Cambridge, UK) for RNA sequencing (Illumina NovaSeq 6000; paired-end 150 bp).

The closed genome sequence of PBIO1953 was annotated with Bakta v. 1.7.0 and bakta database v. 5. In order to process the raw reads and to further quality trim the data, Trim Galore v. 0.6.8 (https://github.com/FelixKrueger/TrimGalore) was used. The trimmed reads were then mapped with Bowtie 2 v. 2.5.1 (mode: –very-sensitive-local), where the assembly of PBIO1953 was used as a reference. The gene counts were calculated using featureCounts v.2.0.1 /stranded-mode) based on PBIO1953śannotation. In the following step, the count table was imported into R v. 4.3.1 (https://www.R-project.org/), and differentially expressed genes were identified with DESeq2 v. 1.40.0 in default mode, with one exception; genes with rowSums of <10 in the count table were excluded before the analysis. Within our analyses, we used an absolute log2 fold change 1.5 threshold combined with an adjusted e value lower than 0.05 to determine differences within gene expression, between the different incubation conditions. We excluded one replicate which was incubated at 37 °C due to a shift on principal component 1 (PC1) and PC2, which can be seen in Table 2 where principal-component analysis is plotted. Finally, all genes were annotated using the online tool eggnog-mapper v. 2.1.9 to extract COG (cluster of orthologous groups) categories.

### Statistical analysis

Statistical analyses were conducted using GraphPad Prism v. 9.5.1 for Windows, developed by GraphPad Software (San Diego, CA US). To assess qPCR results, the Bio-Rad CFX Maestro 2.3 v. 5.3.0 22.1030 (Bio-Rad, Hercules, CA, US) was utilized. All phenotypic experiments consisted of three or more independent biological replicates. Unless otherwise specified, data were presented as the mean and standard deviation. Statistical significance was determined using *p* values below 0.05 to indicate significant differences among the results.

## Author contributions

Conceptualization, E.E. and K.S.; methodology, E.E., L.S., J.U.M and R.S.; software, M.Sc. and S.E.H.; validation, E.E. and J.U.M; formal analysis, E.E., E.A.I., L.S., J.A.B., J.U.M., K.B., K.S., M.Si., M.Sc., R.S., S.G. and S.E.H.; investigation, E.E., J.U.M., M.Si. and R.S.; resources, K.S.; data curation, M.Sc. and S.E.H.; writing—original draft preparation, J.U.M. and E.E.; writing—review and editing, K.S., M.Sc. and S.E.H.; visualization, J.U.M..; supervision, K.S.; project administration, K.S.; funding acquisition, K.S. All authors have read and agreed to the published version of the manuscript.

## Acknowledgments

We thank Sara-Lucia Wawrzyniak for her excellent technical assistance as well as Stefan Bock for excellent technical assistance regarding electron microscopy.

## Data availability

The data for this study have been deposited in the European Nucleotide Archive (ENA) at EMBL-EBI under accession number PRJEB72193 (https://www.ebi.ac.uk/ena/browser/view/PRJEB72193).

## Ethical approval

was given by the ethics committee of the University of Greifswald, Germany (BB 133/20). Informed patient consent was waived as samples were taken under a hospital surveillance framework for routine sampling. The research conformed to the principles of the Helsinki Declaration.

